# Expression of a CO_2_-permeable aquaporin enhances mesophyll conductance in the C_4_ species *Setaria viridis*

**DOI:** 10.1101/2021.04.28.441895

**Authors:** Maria Ermakova, Hannah Osborn, Michael Groszmann, Soumi Bala, Samantha McGaughey, Caitlin Byrt, Hugo Alonso-Cantabrana, Steve Tyerman, Robert T. Furbank, Robert E. Sharwood, Susanne von Caemmerer

## Abstract

A fundamental limitation of photosynthetic carbon fixation is the availability of CO_4_. In C_4_ plants, primary carboxylation occurs in mesophyll cytosol, and little is known about the role of CO_2_ diffusion in facilitating C_4_ photosynthesis. We have examined the expression, localization, and functional role of selected plasma membrane intrinsic aquaporins (PIPs) from *Setaria italica* (foxtail millet) and discovered that *SiPIP2;7 is* CO_2_-permeable. When ectopically expressed in mesophyll cells of *S. viridis* (green foxtail), SiPIP2;7 was localized to the plasma membrane and caused no marked changes in leaf biochemistry. Gas-exchange and C^18^O^16^O discrimination measurements revealed that targeted expression of SiPIP2;7 enhanced the conductance to CO_2_ diffusion from the intercellular airspace to the mesophyll cytosol. Our results demonstrate that mesophyll conductance limits C_4_ photosynthesis at low *p*CO_2_ and that SiPIP2;7 is a functional CO_2_ permeable aquaporin that can improve CO_2_ diffusion at the airspace/mesophyll interface and enhance C_4_ photosynthesis.

---

Diffusion of CO_2_ across biological membranes is a fundamental aspect to photosynthesis. The significant contribution of aquaporins to increased CO_2_ diffusion has been demonstrated in C_3_ plants ^1-3^. Aquaporins have key roles in regulating the movement of water and solutes into roots and between tissues, cells and organelles ^4^. These pore-forming integral membrane proteins can be divided into multiple sub-families depending on their amino acid sequence and sub-cellular localization. The PIPs (plasma membrane intrinsic proteins) are the only sub family, to date, known to permeate CO_2_ ^5^. The PIPs are subdivided into paralog groups PIP1s and PIP2s, based on sequence homology ^6-8^. Typically, PIP2s show higher water permeability when expressed in heterologous systems ^9^ and PIP1s seemingly require interaction with a PIP2 to correctly traffic to the plasma membrane ^10,11^. In plants, a number of CO_2_ permeable PIPs have been identified including *Arabidopsis thaliana* AtPIP1;2 ^12^ and AtPIP2;1 ^13^; *Hordeum vulgare* HvPIP2;1, HvPIP2;2, HvPIP2;3 and HvPIP2;5 ^14^; *Nicotiana tabacum* NtPIP1;5s (NtAQP1) ^15,16^ and *Zea mays* ZmPIP1;5 and ZmPIP1;6 ^17^.

The roles of the CO_2_ permeable aquaporins have been largely characterized in C_3_ photosynthetic plants where aquaporins localized in both the plasma membrane and chloroplast envelopes have been shown to facilitate CO_2_ diffusion from the intercellular airspace to the site of Rubisco in chloroplasts ^18,19^. However, little is known about their role in C_4_ photosynthesis. The C_4_ photosynthetic pathway is a biochemical CO_2_ pump where the initial conversion of CO_2_ to bicarbonate (HCO_3_^-^) by carbonic anhydrase (CA) and subsequent fixation to phospho*enol*pyruvate (PEP) by PEP carboxylase (PEPC) takes place in the cytosol of mesophyll cells. The pathway requires a close collaboration between mesophyll and bundle sheath cells and this constrains leaf anatomy limiting mesophyll surface area that forms a diffusive interface for CO_2_ ^20^. Mesophyll conductance is defined as the conductance to CO_2_ diffusion from the intercellular airspace to the mesophyll cytosol ^21-24^. Although the rate of C_4_ photosynthesis is almost saturated at ambient *p*CO_2_, current modelling suggests that higher mesophyll conductance can increase assimilation rate and water-use-efficiency at low intercellular CO_2_ partial pressures which occur when stomatal conductance is low ^25^.

*Setaria italica* (foxtail millet) and *Setaria viridis* (green foxtail) are C_4_ grasses of the Paniceae tribe and Poaceae family, related to important agronomical crops such as *Z. mays* (maize) and *Sorghum bicolor* (sorghum). *S. viridis* is frequently used as a model species for C_4_ photosynthesis research as it is diploid with a relatively small genome that is sequenced and can be easily transformed ^23,26,27^. Here we used a yeast heterologous expression system to examine the permeability to CO_2_ of selected PIPs from *S. italica*. We identified *SiPIP2;7* as encoding a CO_2_-permeable aquaporin that, when expressed in the plasma membrane of *S. viridis* mesophyll cells, increased mesophyll conductance. Our results demonstrate that CO_2_-permeable aquaporins can be used to increase CO_2_ diffusion from the intercellular airspace to mesophyll cytosol to provide higher carboxylation efficiency in C_4_ leaves.

## Results

### *S. italica* PIP family

Four *PIP1* and eight *PIP2* genes were identified in both *S. italica* and *S. viridis* and their protein sequences were 99–100 % identical between the two species (Table S1). Phylogenetic analysis based on the amino acid sequences of the *S. italica* PIP family showed that three distinct clades emerge: the PIP1 clade, PIP2 clade I, and PIP2 clade II (Fig. S1). Isoforms within these three clades have characteristic differences including sequence signatures associated with substrate selectivity (Table S2). Three of SiPIP1s (1;1, 1;2, 1;5) and all SiPIP2 clade I members (2;1, 2;4, 2;5, 2;6, 2;7) matched the current consensus sequence for CO_2_ transport ^6,28^.

RNA-seq data from the publicly available Phytomine database (Phytozome), was examined for tissue-specific expression patterns of the *S. italica PIPs* (Fig. 1a). *SiPIP1;1, 1;2, 1;5*, and *2;1* express at moderate to high levels and *SiPIP2;6* at low to moderate levels, in all tissues analyzed (root, leaves, shoot, panicle). *SiPIP1;6, 2;4, 2;5, 2;7* and *2;3* were expressed predominantly in roots at low to moderate levels. *SiPIP2;8* was expressed only in leaves and *SiPIP2;2* transcripts were not detected.

**Fig. 1.**
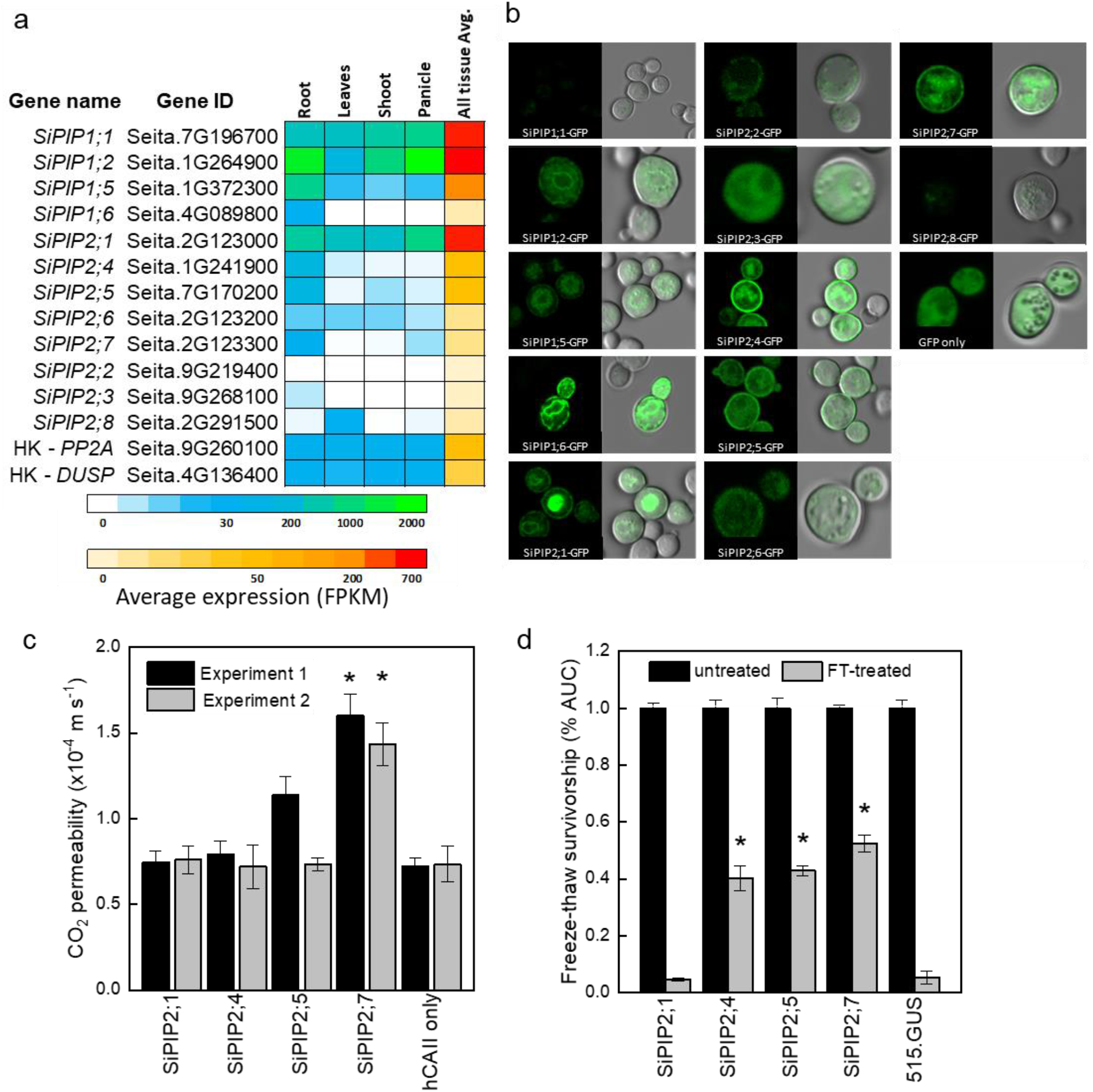
Identification of the CO^2^-permeable aquaporin SiPIP2;7 from *S. italica*. **a**. Expression atlas of the *SiPIP* genes generated from Phytomine reported as Fragments Per Kilobase of transcript per Million mapped reads (FPKM). House-keeping genes (HK) *PROTEIN PHOSPHATASE 2A* (*PP2A*) and *DUAL SPECIFICITY PROTEIN* (*DUSP*) were included for reference. **b**. Localization of SiPIP-GFP fusions expressed in yeast visualised with confocal microscopy; left panels – GFP fluorescence; right panels – bright field overlaid with GFP fluorescence. Measured cell diameters are shown on Fig. S2. **c**. CO^2^ permeability assay on yeast co-expressing *SiPIPs* and *human CARBONIC ANHYDRASE II* (*hCAII*) analyzed by stopped flow spectrometry (see Fig. S2 for details). “hCAII only” expression was used as negative control. Mean ± SE, *n* = 3 biological replicates. Two independent experiments are presented. Asterisks indicate statistically significant differences between yeast expressing *SiPIPs* and “hCAII only” control (*t*-test, *P* < 0.05). **d**. Yeast water permeability assessed in the yeast aquaporin deletion background (*aqy1 aqy2*) by the cumulative growth between untreated and freeze-thawed cells and determined by the percent area under the curve (% AUC). The yeast expressing the β-glucuronidase reporter gene (515.GUS) was used as negative control. Mean ± SE, *n* = 4 biological replicates. Asterisks indicate statistically significant differences between yeast expressing *SiPIPs* and 515.GUS control (*t*-test, *P* < 0.01).

### Functional characterization of PIPs

GFP localization of SiPIP-GFP fusions were used to confirm expression and determine targeting to the yeast plasma membrane (Fig. 1b). Overall, SiPIP1s had lower GFP signal that was patchy at the cell periphery with strong internal signal consistent with localization to the endoplasmic reticulum. GFP signal was also present diffusively throughout the cytosol suggestive of protein degradation. Overall, SiPIP1s were poorly produced in yeast and were not efficiently targeting to the plasma membrane as needed for the functional assays. For the PIP2s, only SiPIP2;1, SiPIP2;4, SiPIP2;5, and SiPIP2;7 showed clear localization to the plasma membrane in addition to other internal structures, and were therefore selected for further functional analyses.

CO_2_ permeability was measured in yeast co-expressing a *SiPIP* along with *human CARBONIC ANHYDRASE II* (*hCAII*). A stopped flow spectrophotometer was used to monitor CO_2_-triggered intracellular acidification via changes in fluorescence intensity of a pH sensitive fluorescein dye Fig. S2; ^12,18,29^. Importantly for reliable results, all SiPIP yeast lines tested showed similar cell volumes and were not limited by CA activity (Fig. S2). A screen of the lines revealed that yeast expressing *SiPIP2;7* had the highest CO_2_ permeability of 1.5 × 10^−4^ m s^-1^, which was significantly larger than the negative control expressing *hCAII* only (Fig. 1c). Other *SiPIP*s displayed comparable CO_2_ permeability to the *hCAII* only control. The changes in CO_2_ permeability detected on the stopped flow spectrophotometer for yeast expressing *SiPIP2;7* were not an artifact brought on by an increased permeability to protons causing the intracellular acidification (Fig. S3).

Freeze-thaw survival assays, which quantify water permeability of aquaporins ^30^, provided further confirmation that the SiPIPs expressed in yeast were functional. Overexpression of water permeable aquaporins greatly improves freeze-thaw tolerance in yeast, especially in the highly compromised aquaporin knockout mutant *aqy1/2* ^30^. Yeast expressing the β-glucuronidase reporter gene (515.GUS) was used a control to show that the single freeze-thaw treatment was effective in almost killing off the entire yeast population (Fig. 1d). Consistent with the poor plasma membrane localization and abundance of SiPIP2;1-GFP (Fig. 1b), yeast expressing *SiPIP2;1* did not show any protection to freeze-thaw treatments (Fig. 1c). On the other hand, *SiPIP2;4, 2;5* and *2;7* all showed some level of protection, indicating that they permeated water and were functional within the plasma membrane of yeast cells. For detailed characterisation of water permeability, SiPIP2;7 was expressed in *Xenopus laevis* oocytes. Swelling assay confirmed that SiPIP2;7 is a functional water channel (Fig. S4).

### Expression of PIP2;7 in mesophyll cells of *S. viridis*

To confirm and exploit the CO_2_ permeability characteristic of SiPIP2;7 *in planta*, we created transgenic *S. viridis* plants expressing *SiPIP2;7* with a C-terminal FLAG-tag fusion and under the control of the mesophyll-preferential *Z. mays* PEPC promoter ^31,32^. Out of 52 T_0_ plants analyzed for SiPIP2;7-FLAG protein abundance and the hygromycin phosphotransferase (*hpt*) gene copy number (Fig. S5), lines 27, 44 and 52 were selected for further analysis because they had the strongest FLAG signal per transgene insertion number. Immunodetection of FLAG and photosynthetic proteins was performed on leaves of homozygous transgenic plants (Fig. 2a); azygous plants of line 44 were used as control hereafter. Monomeric and dimeric SiPIP2;7-FLAG was detected in all transgenic plants (Fig. S5) and abundance of the prevalent dimeric form was used for relative quantification of SiPIP2;7 abundance (Fig. 2a). Plants of line 44 had the highest production of SiPIP2;7-FLAG whilst plants of lines 27 and 52 accumulated about 2-4 times less of this protein. Immunodetection of FLAG on leaf cross-sections, visualized with confocal microscopy, confirmed partial localization of SiPIP2;7-FLAG to the plasma membrane of mesophyll cells (Fig. 2c). Transcript analysis confirmed highly elevated expression of *SiPIP2;7-FLAG* in leaves, but not in roots of transgenic lines (Fig. S6).

**Fig. 2.**
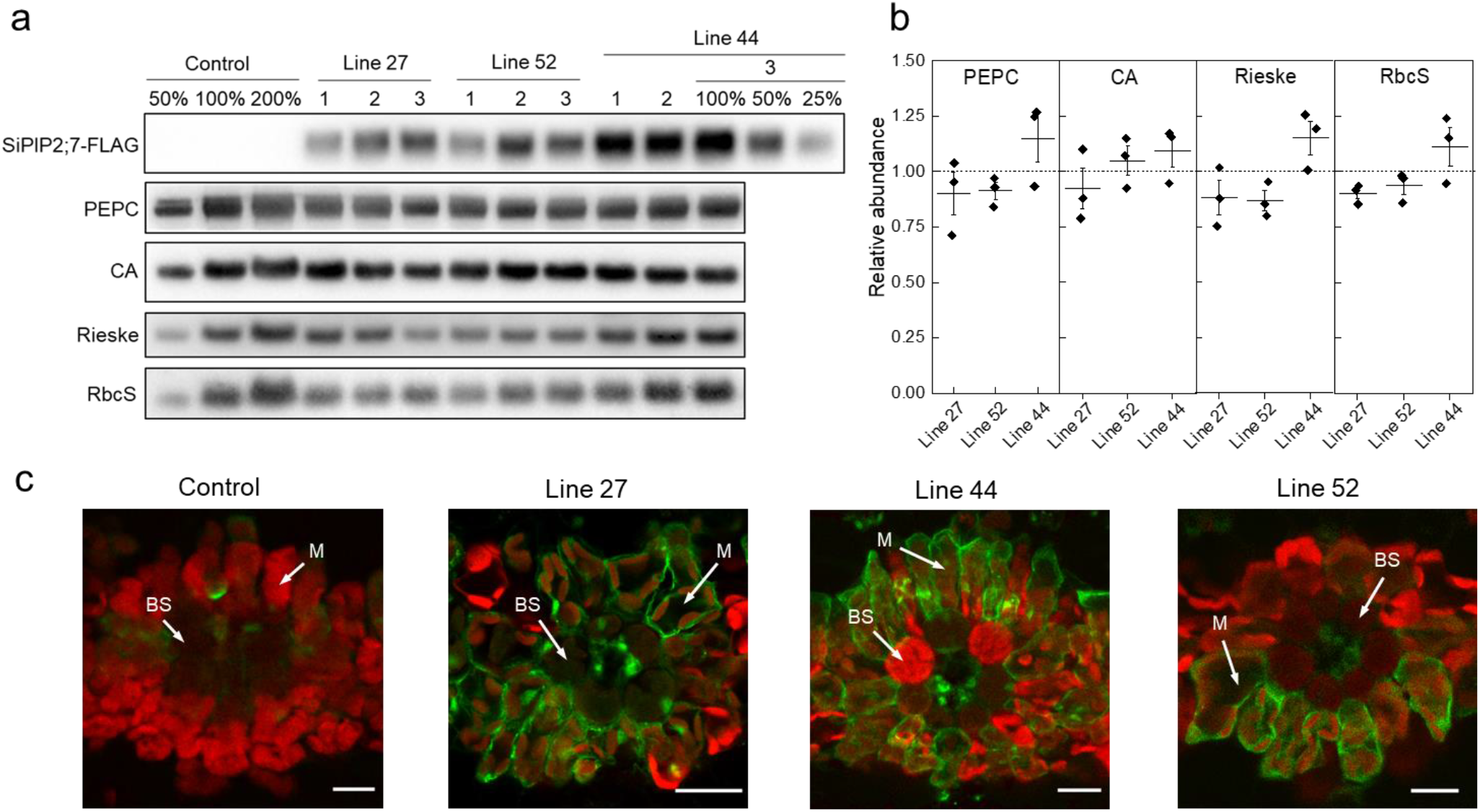
Characterization of *S. viridis* plants expressing *SiPIP2;7-FLAG* in mesophyll cells. **a**. Immunodetection of SiPIP2;7-FLAG and photosynthetic proteins in leaf protein samples loaded on leaf area basis. Three plants from each of the three transgenic lines were analyzed and dilution series of the control and line 44-3 samples were used for relative quantification. **b**. Protein abundances calculated from the immunoblots relative to control plants. Mean ± SE. No significant difference was found between the transgenic and control plants (*t*-test, *P* < 0.05). **c**. Immunolocalisation of SiPIP2;7-FLAG on leaf cross-sections visualized with confocal microscopy. Fluorescence signals are pseudo-colored: green -FLAG antibodies labelled with secondary antibodies conjugated with Alexa Fluor 488; red -chlorophyll autofluorescence. BS, bundle sheath cell; M, mesophyll cell. Scale bars = 20 µm. Azygous plants of line 44 were used as control.

Abundances of photosynthetic proteins PEPC, CA, the Rieske subunit of the Cytochrome *b*_6_ *f* complex, and the small subunit of Rubisco (RbcS), did not differ between transgenic and control plants (Fig. 2a). In line with the immunoblotting results, measured activities of PEPC and CA, and the amount of Rubisco active sites were not altered in the transgenic plants (Table 1). Chlorophyll content, leaf dry weight per area and biomass of roots and shoots did not differ between the genotypes either (Table 1).

**Table 1.** Properties of *S. viridis* plants expressing *SiPIP2;7-FLAG* in mesophyll cells. PEPC, PEP carboxylase; Rubisco, ribulose bisphosphate carboxylase oxygenase; LMA, leaf mass per area. Azygous plants of line 44 were used as control. Mean ± SE, *n* = 3 except for biomass (*n* = 8). Three-weeks old plants before flowering were used for all analyses. No significant difference was found between the transgenic and control plants (One-way ANOVA, α = 0.05).

| Parameter | Control | Line 27 | Line 44 | Line 52 |
| --- | --- | --- | --- | --- |
| PEPC activity, $\mu\text{mol CO}_2 \text{ m}^{-2} \text{ s}^{-1}$ | 220.1 ±25.8 | 197.6 ±12.7 | 208.7 ±7.9 | 218.5 ±3.5 |
| CA hydration rate, $\text{mol m}^{-2} \text{ s}^{-1} \text{ bar}^{-1}$ | 6.50 ±0.10 | 6.32 ±0.22 | 5.34 ±0.67 | 5.35 ±0.56 |
| Rubisco active sites, $\mu\text{mol m}^{-2}$ | 12.17 ±0.63 | 12.53 ±0.54 | 12.84 ±0.13 | 12.63 ±0.74 |
| Chlorophyll (*a+b*), $\text{mmol m}^{-2}$ | 0.71 ±0.07 | 0.72 ±0.04 | 0.72 ±0.05 | 0.72 ±0.08 |
| Chlorophyll *a/b* | 5.01 ±0.16 | 5.08 ±0.05 | 4.97 ±0.09 | 5.07 ±0.15 |
| LMA, g (dry weight) $\text{m}^{-2}$ | 23.6 ±1.6 | 24.0 ±1.5 | 25.6 ±1.3 | 25.4 ±1.3 |
| Shoot biomass, g (dry weight) $\text{plant}^{-1}$ | 2.06 ±0.36 | 2.01 ±0.20 | 2.23 ±0.31 | 2.24 ±0.34 |
| Root biomass, g (dry weight) $\text{plant}^{-1}$ | 0.27 ±0.07 | 0.28 ±0.03 | 0.34 ±0.06 | 0.35 ±0.05 |

To study the effects of *SiPIP2;7-FLAG* ectopic expression on photosynthetic properties in the transgenic plants, we conducted concurrent gas-exchange and fluorescence analyses at different intercellular CO_2_ partial pressure (*C*_i_) (Fig. 3). No significant changes were detected between transgenic and control plants in CO_2_ assimilation rates (*A*), effective quantum yield of Photosystem II (φPSII) or stomatal conductance to water vapor at ambient CO_2_ (Fig. S7). However, since CO_2_ assimilation rates were consistently higher in all transgenic plants at low *C*_i_ (Fig. 3a, inset), we analyzed the initial slopes of the CO_2_ response curves and mesophyll conductance. Fitting linear regressions (Fig. 4a) indicated that lines 44 and 52 had significantly greater initial slopes (average values of 0.52 and 0.53, respectively) compared to the control (0.41), whereas line 27 had a slightly increased initial slope (0.46).

**Fig. 3.**
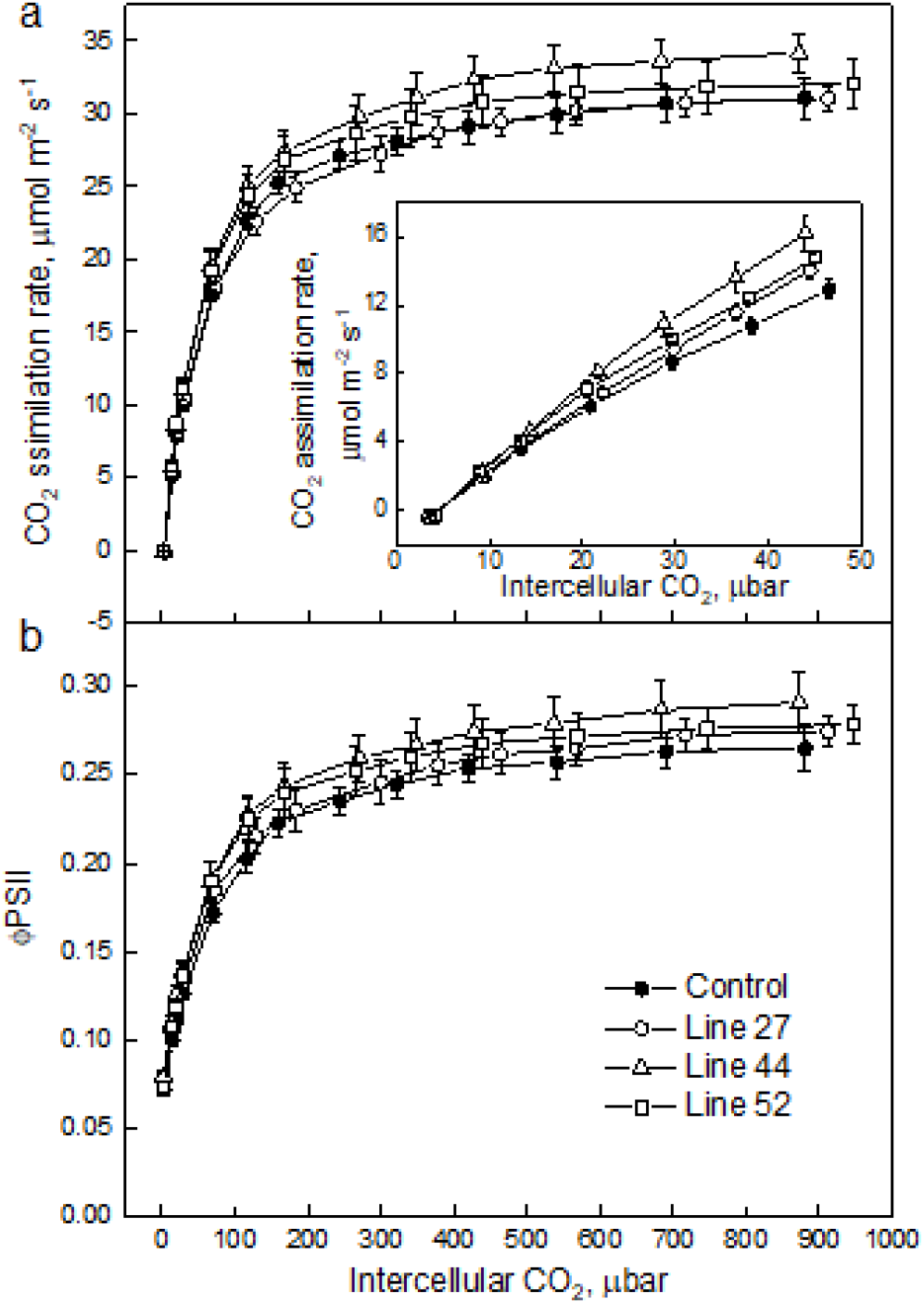
CO^2^ response of CO^2^ assimilation rate (**a**) and quantum yield of Photosystem II (**b**) in *S. viridis* plants expressing *SiPIP2;7-FLAG* in mesophyll cells. Measurements were performed at the irradiance of 1500 µmol m^-2^ s^-1^; azygous plants of line 44 were used as control. Mean ± SE, *n* = 4-5 biological replicates. No significant difference was found between the transgenic and control plants (One-way ANOVA, α = 0.05).

**Fig. 4.**
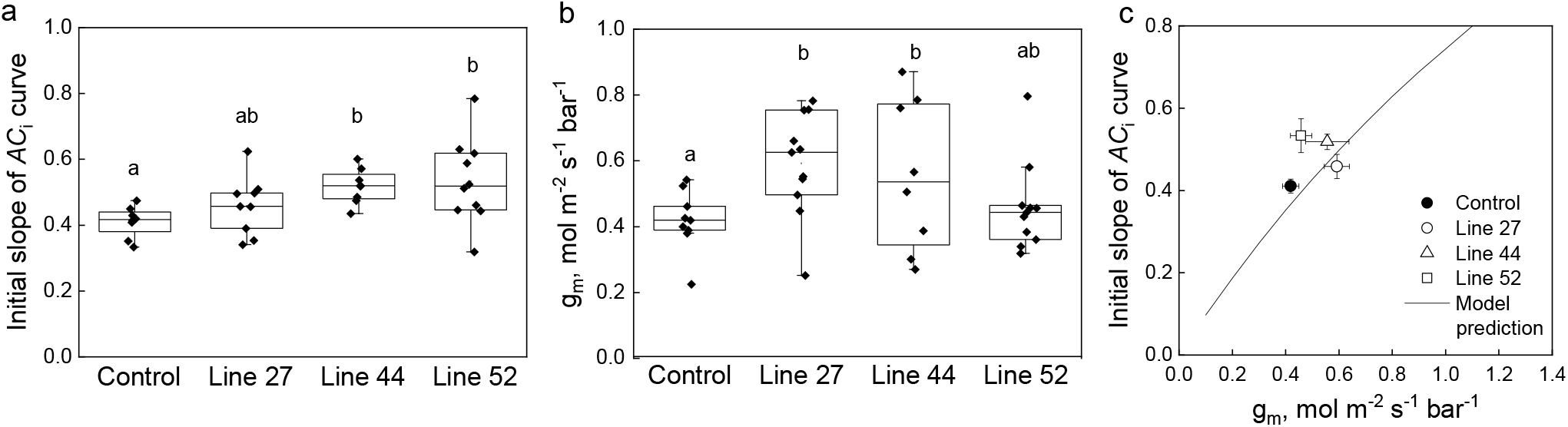
Effect of the mesophyll conductance, *g*^m,^on the initial slope of the CO^2^ assimilation response curve to the intercellular CO^2^ partial pressure (*AC*^i^ curve) in leaves of *S. viridis* expressing *SiPIP2;7-FLAG* in mesophyll cells. **a**. Mesophyll conductance, *g*^m^, estimated by oxygen isotope discrimination assuming full isotopic equilibrium ^23^. Measurements were made at ambient CO^2^ and low O^2^. **b**. Initial slope of the *AC*^i^ curves estimated by linear fitting of curves presented in Fig. 3a inset. **c**. Data from a and b compared to the C^4^ biochemical model predictions ^36^. The model relates the initial slope of the *AC*^i^ curve (d*A*/*C*^i^) to *g*^m,^ by: 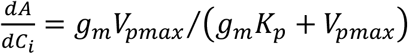, where *V* _pmax_ and *K* _p_ denote the maximum PEPC activity and the Michaelis Menten constant for CO2 taken here as 250 µmol m^-2^ s^-1^ and 82 µbar ^65,66^. Azygous plants of line 44 were used as control. Letters indicate statistically significant differences between the groups (One-way ANOVA with Tukey post-hoc test, α = 0.05).

### Mesophyll conductance to CO_2_ in plants expressing SiPIP2;7

Measurements of Δ^18^O were used to estimate conductance of CO_2_ from the intercellular airspace to the sites of CO_2_ and H_2_O exchange in the mesophyll cytosol (*g*_m_) with the assumption that CO_2_ was in full isotopic equilibrium with leaf water in the cytosol ^23,33^. Transgenic lines 27 and 44 had significantly greater mesophyll conductance than control plants (0.42 mol m^-2^ s^-1^ bar^-1^) with average values of 0.59 and 0.55 mol m^-2^ s^-1^ bar^-1^, respectively (Fig. 4b). We also used the *g*_m_ calculations proposed by Ogée *et al*. ^34^ which try to account for the rates of bicarbonate consumption by CA. The CA hydration constant (*k*_CA_) of 6.5 mol m^-2^ s^-1^ bar^-1^ was used for these calculations (Table 1). We found that the *g*_m_ measured with this method gave on average 1.25 times greater values but did not change the ranking of mesophyll conductance shown in Fig. 4a (Fig. S8). The C_4_ photosynthetic model by von Caemmerer and Furbank ^35^ and von Caemmerer ^36^ relates the initial slope of the CO_2_ response curve (d*A*/*C*_i_) to *g*_m_ (see Fig. 4 caption and Materials and Methods). Fig. 4c shows that the measured relationship between the initial slope and *g*_m_ fits closely with model prediction.

## Discussion

The diffusion of CO_2_ from the earth’s atmosphere to the site of primary carboxylation within leaves of C_3_ and C_4_ plants often limits photosynthesis and impacts the efficient use of water. In C_4_ plants, primary carboxylation occurs in mesophyll cytosol and a large mesophyll conductance, *g*_m_, is required to account for high photosynthetic rates which generate a large drawdown between the intercellular airspace and the cytosol ^21^. An effective strategy to enhance CO_2_ diffusion in C_3_ plants has been the overexpression of CO_2_ permeable aquaporins in plasma membrane and the chloroplast envelope leading to improved *g*_m_, assimilation rate or grain yield ^1,3,15,37^. Screening *S. italica* PIPs for CO_2_ permeability in a yeast heterologous system resulted in identification of SiPIP2;7 as a CO_2_ pore (Fig. 1c). Expression analysis revealed that *SiPIP2;7* was almost exclusively expressed in roots under ideal conditions (Fig. 1a, Fig. S6) which, combined with the water permeability identified in yeast and oocyte assays (Fig. 1d, Fig. S4), suggest that SiPIP2;7 may function in regulating root hydraulic conductivity, a role extensively documented for PIP aquaporins ^38,39^. The physiological relevance of SiPIP2;7’s CO_2_ permeating capacity is not immediately clear. Gas uptake by roots is well documented ^40^ and in C_3_ plants CO_2_ uptake by roots may contribute to the C_4_ photosynthesis-like metabolism detected in stems and petioles ^41^. It is possible that *SiPIP2;7* is conditionally expressed in leaves, or even that its capacity to transport CO_2_ is inadvertent and related to the transportation of another yet undetermined substrate; analogous to the uptake of toxic metalloids by some NIP aquaporins due to their capacity to transport boron ^42^. Further work is needed to determine whether PIPs in general function natively as relevant CO_2_ pores in C_4_ leaves.

We employed the CO_2_ transport capacity of SiPIP2;7 to enhance transmembrane CO_2_ diffusion from the intercellular airspace into the mesophyll cytosol, where CA and PEPC reside, by overexpressing *SiPIP2;7* in *S. viridis*. We confirmed the localization of SiPIP2;7 within the mesophyll plasma membranes (Fig. 2c) and detected the increase in CO_2_ diffusion across the mesophyll membranes in transgenic plants by two independent methods. First, we calculated *g*_m_ from the C^18^O^16^O discrimination measurements (Fig. 4b) and the theory for these calculations has been outlined ^23,33,43^. Second, we fitted linear regressions to the initial slopes of the *AC*_i_ curves (Fig. 3a inset, Fig. 4a), which depend on *g*_m_, *V*_pmax_ and *K*_p_ where the two latter parameters denote the maximum PEPC activity and the Michaelis Menten constant of PEPC for HCO_3_ ^-35,36^. Since PEPC and CA activities were not altered in plants expressing *SiPIP2;7* (Table 1), higher initial slopes of the *AC*_i_ curves in transgenic lines were attributed to the increased *g*_m_. Up-regulation of *g*_m_ in lines 27 and 52 was confirmed by one of the methods, while both methods indicated significantly increased *g*_m_ in line 44 (Fig. 4). When plotted against each other, the initial slopes and *g*_m_ in transgenic and control plants, fitted the model predictions confirming the hypothesised functional role of *g*_m_ in C_4_ photosynthesis ^24,36,44^. Our findings explicitly demonstrate that mesophyll conductance limits C_4_ photosynthesis at low CO_2_ and indicate that increasing CO_2_ diffusion at the airspace/mesophyll interface, combined with complementary traits including overexpression of Cytochrome *b*_6_ *f* and Rubisco ^27,31^, could further improve C_4_ photosynthesis.

## Materials and methods

### Heterologous expression in yeast

cDNAs encoding the 12 *S. italica* aquaporins (Table S1) and *human CARBONIC ANHYDRASE II* (AK312978) were codon-optimized for expression in yeast with IDT DNA tool (https://sg.idtdna.com/pages/tools) and a yeast related kozak sequence was added at the 5’ end to help increase translation ^45^. For CO_2_ permeability measurements, pSF-TPI1-URA3 with an aquaporin and pSF-TEF1-LEU2 with hCAII were co-transformed into the *S. cerevisiae* strain INV*Sc*1 (Thermo Fisher Scientific, Waltham, MA). For water permeability measurements, pSF-TPI1-URA3 with an aquaporin was transformed into the *aqy1/2* double mutant yeast strain deficient in aquaporins ^46^. The yeast vectors pSF-TPI1-URA3 and pSF-TEF1-LEU2 were obtained from Oxford Genetics (Oxford, UK). Yeast transformation was performed using the Frozen-EZ yeast transformation II kit (Zymo Research, Irvine, CA) and selection of positive transformants was based on amino acid complementation. To ensure CA was not limiting, CA activity was determined using a membrane inlet mass spectrometry as described by Endeward, et al. ^47^ (Fig. S2). For CO_2_ permeability measurements an average cell diameter of 4.63 µm was determined by measuring ∼100 yeast cells expressing each aquaporin (Fig. S2). To study the subcellular localizations of aquaporins in yeast, a C-terminus GFP tag was added to the sequences into the pSF-TPI1-URA3 vector (pSF-TPI1-URA3-GFP). The fluorescence signal was observed using a Zeiss 780 confocal laser scanning microscope (Zeiss, Oberkochen, Germany): excitation 488 nm, emission 530 nm. Cytosolic GFP expression was used as control.

### CO_2_ induced intracellular acidification assay

CO_2_ intracellular acidification was measured in yeast cells loaded with fluorescein diacetate (Sigma-Aldrich, St. Louis, MO) as described previously ^48,49^. Briefly, an overnight culture of yeast cells was collected and resuspended in an equal volume of 50 mM 4-(2-hydroxyethyl)-1-piperazineethanesulfonic acid (HEPES)-NaOH, pH 7.0, 50 µM fluorescein diacetate and incubated for 30 min in the dark at 37 °C. The suspension was centrifuged and the pellet resuspended in ice-cold incubation buffer (25 mM HEPES-NaOH, pH 6.0, 75 mM NaCl). Cells loaded with fluorescein diacetate were then injected into the stopped flow spectrophotometer (DX.17MV; Applied Photophysics, Leatherhead, UK) alongside a buffer solution (25 mM HEPES, pH 6.0, 75 mM NaHCO_3_, bubbled with CO_2_ for 2 h). The kinetics of acidification was measured at 490 nm excitation and >515 nm emission (OG515 long pass filter, Schott, supplied by Applied Photophysics). Data was collected over a time interval of 0.2 s and analysed using ProData SX viewer software (Applied Photophysics). CO_2_ permeability was determined using the method of Yang, et al. ^50^. An average of 75 injections over at least three separate cultures was used for each aquaporin.

### Determination of water permeability

A freeze-thaw yeast assay was used to determine water permeability of aquaporins expressed in *aqy1/2* based on previous reports ^30^. Briefly, an overnight culture was diluted to ∼6×10^6^ cells (final volume 1 mL) in appropriate selection liquid growth medium and incubated at 30 °C for 1 h. 250 µL of each culture were then aliquoted into two standard 1.5 mL microtubes: the first (control) tube was placed on ice and the second tube was subject to a single freeze-thaw treatment, consisting of 30-s freezing in liquid nitrogen and thawing for 20 min in a 30 °C water bath. Following the treatment, the cells were placed on ice. The tubes were then vortexed briefly to ensure even suspension of cells and 200 µL of the culture was transferred to wells of a Nunc-96 400 µL flat bottom untreated plate (Thermo Fisher Scientific, Cat#243656). Yeast growth in control and treated cultures were monitored over a 24-30 h period in a M1000 Pro plate reader (TECAN, Männedorf, Switzerland) at 30 °C with double orbital shaking at 400 rpm and measuring absorbance at 650 nm every 10 min. Growth data was log transformed and freeze-thaw survival calculated as the growth (area under the curve) of treated culture relative to its untreated control from time zero up until the untreated control culture reached stationary phase.

For swelling assays, the coding sequence of *SiPIP2;7* was cloned into pGEMHE oocyte expression vector using LR clonase II (Thermo Fisher Scientific) and cRNA was synthesised with mMessage mMachine® T7 Transcription Kit (Thermo Fisher Scientific). *Xenopus laevis* oocytes were injected with 46 nL of RNAse-free water with either no cRNA or 23 ng cRNA with a micro-injector Nanoinject II (Drummond Scientific, Broomall, PA). Post-injection oocytes were stored at 18 °C in a Low Na^+^ Ringer’s solution [62 mM NaCl, 36 mM KCl, 5 mM MgCl_2_, 0.6 mM CaCl_2_, 5 mM HEPES, 5% (v/v) horse serum (H-1270, Sigma-Aldrich) and antibiotics: 0.05 mg mL^-1^ tetracycline, 100 units mL^-1^ penicillin/0.1 mg mL^-1^ streptomycin], pH 7.6 for 24–30 h. Photometric swelling assay was performed 24-30 h post-injection ^51^.

### Construct assembly and *S. viridis* transformation

The coding sequence of *S. viridis PIP2;7* (Sevir.2G128300.1, Phytozome, https://phytozome.jgi.doe.gov/) has been codon optimized for the Golden Gate cloning ^52^ and translationally fused with the glycine linker and the FLAG-tag coding sequence ^53^. The resulting coding sequence was assembled with the *Z. mays PEPC* promoter and the bacterial tNos terminator into the second expression module of the pAGM4723 binary vector. The first expression module has been occupied by the hygromycin phosphotransferase (*hpt*) gene assembled with the *Oryza sativa* actin promoter and the tNos terminator. The construct was transformed into *S. viridis* cv. MEO V34-1 using *Agrobacterium tumefaciens* strain *AGL1* following the procedure described in Osborn, et al. ^23^. T_0_ plants resistant to hygromycin were transferred to soil and analyzed for *hpt* insertion number by droplet digital PCR (iDNA Genetics, Norwich, UK). The T_1_ and T_2_ progenies of T_0_ plants 27, 44 and 52 were analyzed. Azygous T_1_ plants of line 44 and their progeny were used as control.

### Plant growth conditions

Seeds were surface-sterilized and germinated on medium (pH 5.7) containing 2.15 g L^-1^ Murashige and Skoog salts, 10 mL L^-1^ 100x Murashige and Skoog vitamins stock, 30 g L^-1^ sucrose, 7 g L^-1^ Phytoblend, 20 mg L^-1^ hygromycin (no hygromycin for azygous plants). Seedlings that developed secondary roots were transferred to 0.6 L pots with garden soil mix layered on top with 2 cm seed raising mix (Debco, Tyabb, Australia) both containing 1 g L^-1^ Osmocote (Scotts, Bella Vista, Australia). Plants were grown in controlled environmental chambers with 16 h light/8 h dark, 28 °C day, 22 °C night, 60% humidity and ambient CO_2_ concentrations. Light intensity of 300 µmol m^-2^ s^-1^ was supplied by 1000 W red sunrise 3200 K lamps (Sunmaster Growlamps, Solon, OH). Youngest fully expanded leaves of the 3–4 weeks plants before flowering were used for all analyses.

### Chlorophyll and enzyme activity

Chlorophyll content was measured on frozen leaf discs homogenised with a TissueLyser II (Qiagen, Venlo, The Netherlands) ^54^. PEPC activity was determined after Pengelly, et al. ^55^ from fresh leaf extracts from the plants adapted for 1 h to 800 µmol photons m^-2^ s^-1^. CA activity was measured on a membrane inlet mass spectrometer as a rate of ^18^O exchange from labelled ^13^C^18^O_2_ to H_2_ ^16^O at 25 °C according to von Caemmerer, et al. ^56^ by calculating the hydration rate after Jenkins, et al. ^57^. The amount of Rubisco active sites was determined by [^14^C] carboxyarabinitol bisphosphate binding as described earlier ^58^.

### RNA isolation and qPCR

Leaf and root tissue were frozen in liquid N_2_. Leaf samples were homogenised using a TissueLyser II and RNA was extracted using the RNeasy Plant Mini Kit (Qiagen). Roots were ground with mortar and pestle in liquid N_2_ and RNA was isolated according to Massey ^59^. Briefly, 150 µL of pre-heated (60 °C) extraction buffer [0.1 M trisaminomethane (Tris)-HCl, pH 8, 5 mM ethylenediaminetetraacetic acid (EDTA), 0.1 M NaCl, 0.5% sodium dodecyl sulfate (SDS), 1% 2-mercaptoethanol) was added to ∼100 mg of fine root powder and incubated at 60 °C for 5 min. 150 µL of phenol:chloroform:isoamyl alcohol (25:24:1) saturated with 10 mM Tris (pH 8.0) and 1 mM EDTA was added to the samples, vortexed vigorously for 10 min and centrifuged at 4500 g for 15 min. Aqueous phase was mixed with 120 µL of isopropanol and 15 µL of 3 M sodium acetate and incubated at -80 °C for 15 min, then centrifuged at 4500 g (30 min, 4 °C). The pellet was washed twice in 300 µL of ice-cold 70% ethanol, air dried and dissolved in 60 µL of RNase-free water. After addition of 40 µL of 8 M LiCl, samples were incubated overnight at 4 °C. Nucleic acids were pelleted by centrifugation at 16,000 g (60 min, 4 °C), washed twice with 200 µL of ice-cold 70% ethanol, air dried and dissolved in RNase-free water. DNA from the samples was removed using an Ambion TURBO DNA free kit (Thermo Fisher Scientific), and RNA quality was determined using a NanoDrop (Thermo Fisher Scientific). 100 ng of total RNA were reverse transcribed into cDNA using a SuperScript™III Reverse Transcriptase (Thermo Fisher Scientific). qPCR and melt curve analysis were performed on a Viia7 Real-time PCR system (Thermo Fisher Scientific) using the Power SYBR green PCR Master Mix (Thermo Fisher Scientific) according to the manufacturer’s protocol.

Primer pairs designed to distinguish between *S. viridis PIP2;6* and *PIP2;7* using Primer3 in Geneious Prime (https://www.geneious.com) and reference primers are listed in Table S3.

### Western blotting and immunolocalization

Protein isolation from leaves and gel electrophoresis were performed as described earlier ^27^. Proteins were probed with antibodies against FLAG (ab49763, 1:5000, Abcam, Cambridge, UK), RbcS ^60^ (1:10,000), Rieske (AS08 330, 1:3000, Agrisera, Vännäs Sweden), PEPC (AS09 458, 1:10,000, Agrisera), CA ^61^ (1:10,000). Quantification of immunoblots was performed with Image Lab software (Biorad, Hercules, CA). For immunolocalization leaf tissue was fixed and probed with primary antibodies against FLAG (1:40) and secondary goat anti-mouse Alexa Fluor 488-conjugated antibodies (ab150113, 1:200, Abcam) as described in Ermakova, et al. ^62^. Images were captured with a Zeiss 780 microscope using ZEN 2012 software (Black edition, Zeiss, Oberkochen, Germany). Images for plants of lines 27, 44 and azygous plants were acquired using online fingerprinting (488 nm excitation) with three user-defined spectral profiles for AlexaFluor488, endogenous autofluorescence and chlorophyll. The spectral profile for endogenous autofluorescence was derived from the azygous control. The image for line 52 was initially collected as a full spectral scan (490-660 nm), then linearly un-mixed using the same online fingerprint settings as previously described. Images were post-processed with FIJI ^63^, and histograms for all images were min-max adjusted.

### Gas exchange measurements

Gas-exchange and fluorescence analysis were performed at an irradiance of 1500 µmol m^-2^ s^-1^ (90% red/10% blue actinic light) and different intercellular CO_2_ partial pressures using a LI-6800 (LI-COR Biosciences, Lincoln, NE) equipped with a fluorometer head 6800-01 A (LI-COR Biosciences). Leaves were first equilibrated at 400 ppm CO_2_ in the reference side, leaf temperature 25 °C, 60% humidity and flow rate 500 µmol s^-1^ and then a stepwise increase of CO_2_ concentrations from 0 to 1600 ppm was imposed at 3 min intervals. Initial slopes of the CO_2_ response curves were determined by linear fitting in OriginPro 2018b (OriginLab, Northampton, MA). Quantum yield of PSII upon the application of multiphase saturating pulses (8000 µmol m^-2^ s^-1^) was calculated according to Genty, et al. ^64^.

### C^18^O^16^O discrimination measurements

Simultaneous measurements of exchange of CO_2_, H_2_O, C^18^O^16^O and H_2_ ^18^O were made by coupling two LI-6400XT gas-exchange systems to a tunable diode laser (TDL: model TGA200A, Campbell Scientific Inc., Logan, UT) to measure C^18^O^16^O discrimination and a Cavity Ring-Down Spectrometer (L2130-i, Picarro Inc., Sunnyvale, CA) to measure the oxygen isotope composition of water vapor ^23^. Measurements were made at 2% O^2^, 380 µmol mol^-1^ CO^2^, leaf temperature of 25 °C, irradiance of 1500 µmol m^-2^ s^-1^ and relative humidity of 55%. Each leaf was measured at 4 min intervals and 10 readings were taken. Mesophyll conductance was calculated as described by Osborn, et al. ^23^ with the assumptions that there was sufficient carbonic anhydrase (CA) in the mesophyll cytosol for isotopic equilibration between CO^2^ and HCO_3_ ^-^ We also used the calculations proposed by Ogée, et al. ^34^ to estimate *g*^m.^These calculations try to account for the rates of bicarbonate consumption by CA. We used the rate constant of CA hydration (*k*^CA^) of 6.5 mol m^-2^ s^-1^ bar^-1^ for these calculations.

## Statistical analysis

One-way ANOVAs with Tukey post-hoc test were performed in OriginPro 2018b. A two-tailed, heteroscedastic Student’s *t*-tests were performed in Microsoft Excel.

## Data availability

The datasets and materials generated during the current study are available from the corresponding authors on request.

## The authors declare no competing interests

## Acknowledgements and funding sources

We thank Xueqin Wang for *S. viridis* transformation, Zac Taylor for gas-exchange measurements, Murray Badger and Dimitri Tolleter for measuring CA activity in yeast, Daryl Webb, Ayla Manwaring and the Centre for Advanced Microscopy at the Australian National University for confocal imaging, Wendy Sullivan for help with the stopped flow spectrophotometry and Nerea Ubierna for sharing her spreadsheet for the Ogee *et al. g*^m^calculations. Funding information: this research was supported by the Australian Research Council (ARC) Centre of Excellence for Translational Photosynthesis (CE140100015). RES was funded by ARC DECRA (DE130101760). This work is presented in the Australian provisional patent application # 2021900409.

## Author contributions

RES, SVC, RTF and ME designed the research. ME, HO, MG, SB, SM, RES and SVC performed experiments. ME, RES, SVC and HO wrote the manuscript with contribution of MG. All authors contributed to data analysis and manuscript editing.

