## Supplementary information for "Expression of a CO_2_-permeable aquaporin enhances mesophyll conductance in the C_4_ species *Setaria viridis*"

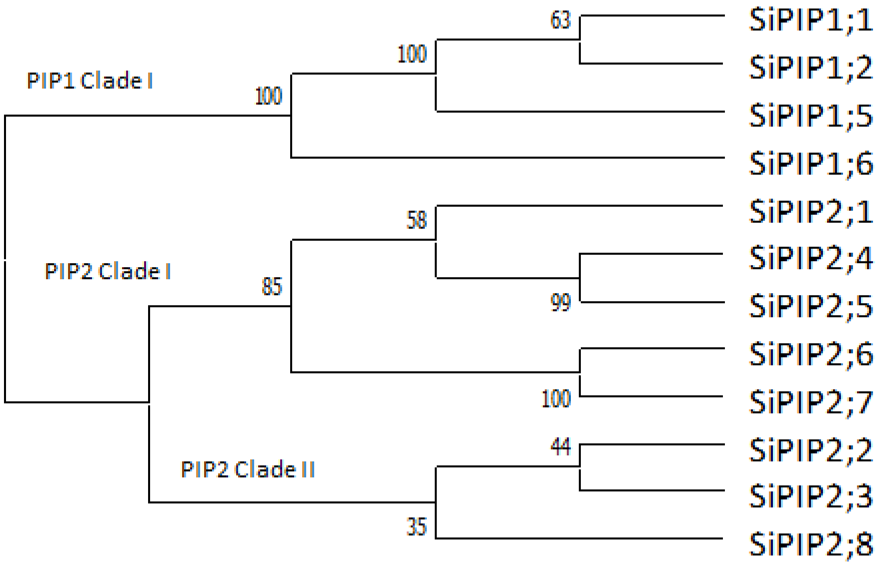


**Fig. S1**. Phylogenetic analysis of *S. italica* PIP protein sequences. Amino acid sequences for the PIP1 and PIP2 proteins from *Setaria italica* were acquired from Phytozome and numbered based on McGaughey, et al. [67]. The protein sequences were aligned using MUSCLE in Geneious (v 11.1.5) and the phylogenetic tree generated using the neighbour-joining method with pairwise deletions in MEGA10 [68].

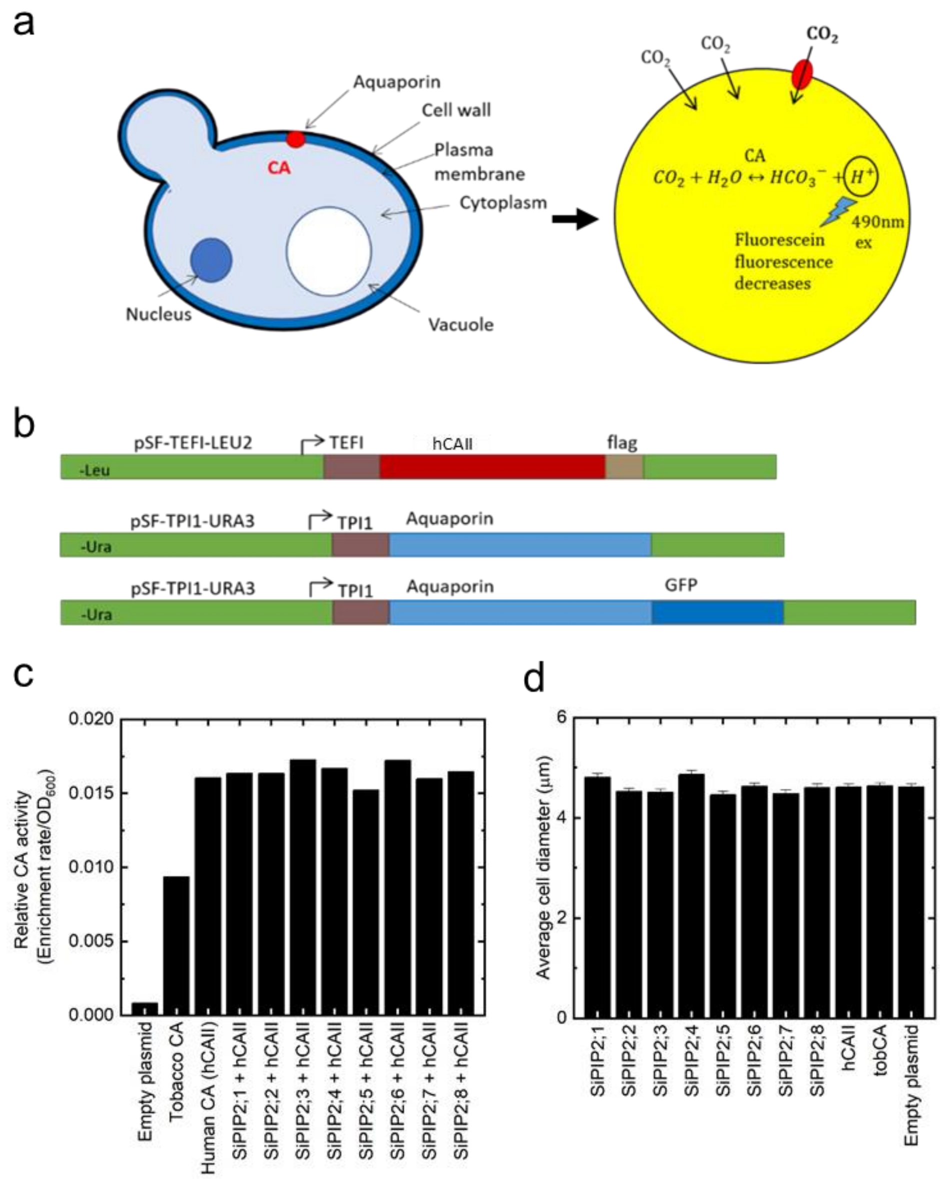


**Fig. S2**. Expression of *S. italica* PIPs in yeast for functional testing of $CO_2$ permeability. **a**. Schematic highlighting the yeast expression system and the detection of $CO_2$ permeability by measuring the change in fluorescein fluorescence due to the acidification of the cytoplasm by producing protons from the hydration of $CO_2$ from ectopically expressed carbonic anhydrase. The decline in fluorescence is measured by stop-flow spectrophotometry at 490nm. **b**. Construct design used for expression and assay of $CO_2$ permeability and localisation in yeast. **c.** CA activity measured in yeast cells expressing PIPs. **d**. The cell diameter of yeast cells expressing PIPs.

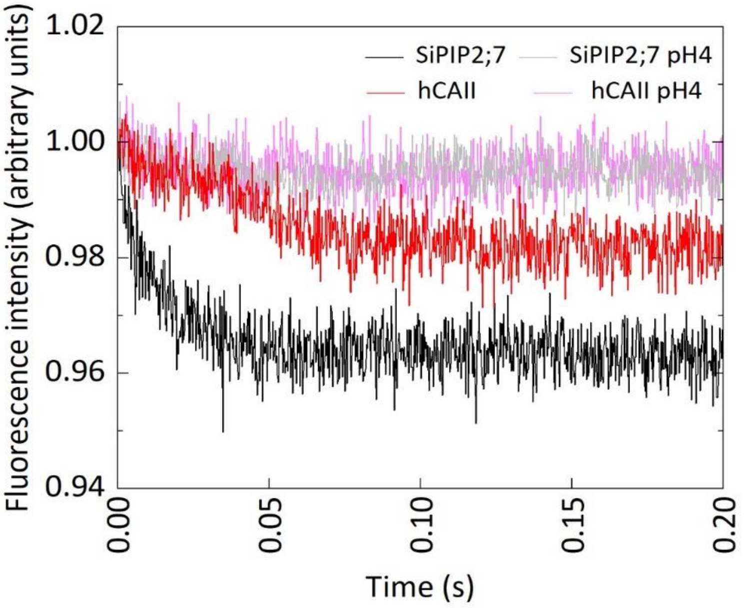


**Fig. S3.** $CO_2$ transport capacity of SiPIP2;7 is not confounded by permeability of protons. The changes in $CO_2$ permeability, detected on the stopped flow spectrophotometer (Fig. 1c), could be confounded by other factors such as permeability to protons also causing intracellular acidification. When SiPIP2;7 + hCAII (black) and hCAII alone (red) where injected alongside $CO_2$ enriched buffer, a decrease in pH was observed. This decrease was more pronounced in PIP2;7 + hCAII indicating higher $CO_2$ transport rate. When SiPIP2;7 + hCAII and hCAII were injected with a low pH (*i.e*. proton rich) buffer, no decrease in fluorescence intensity was observed with either SiPIP2;7 + hCAII (grey) or hCAII alone (pink) demonstrating that the decrease in fluorescence seen with the $CO_2$ enriched was not due to proton movement through the membrane. In fact, both traces were the same specifically pointing out that SiPIP2;7 expression did not cause excessive proton movement which could lead to a false-positive result for $CO_2$ permeability.

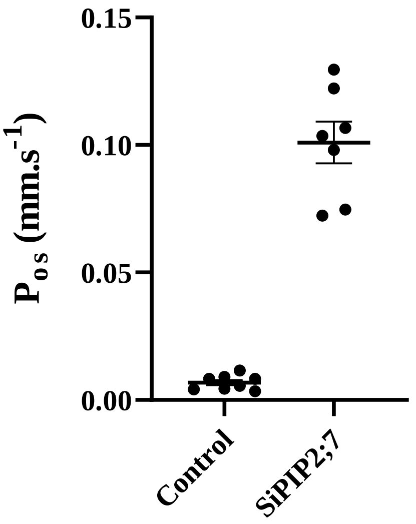


**Fig. S4**. SiPIP2;7 is a functional water channel when expressed in *X. laevis* oocytes. Oocytes were injected with either water (control) or 23 ng of *SiPIP2;7* cRNA. Osmotic permeability ($P_{os}$) was determined via photometric swelling where oocytes were pre-incubated in the isotonic swelling solution before being transferred to the hypotonic swelling solution for the assay. Mean ± SE, $P < 0.0001$ (*t*-test).

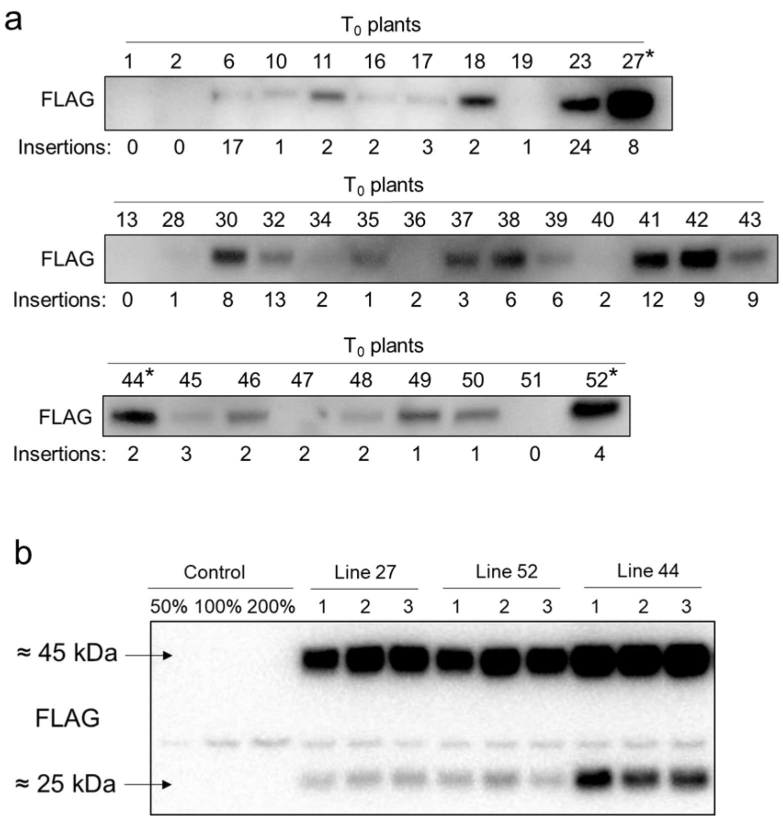


**Fig. S5**. Immunodetection of SiPIP2;7-FLAG in leaves of transgenic *S. viridis* plants. **a**. $T_0$ plants; T-DNA insertion numbers indicate the number of *hpt* gene copies detected by droplet digital PCR. Lines selected for further analysis are marked with asterisks. **b**. Homozygous $T_3$ plants; azygous plants of line 44 were used as control.

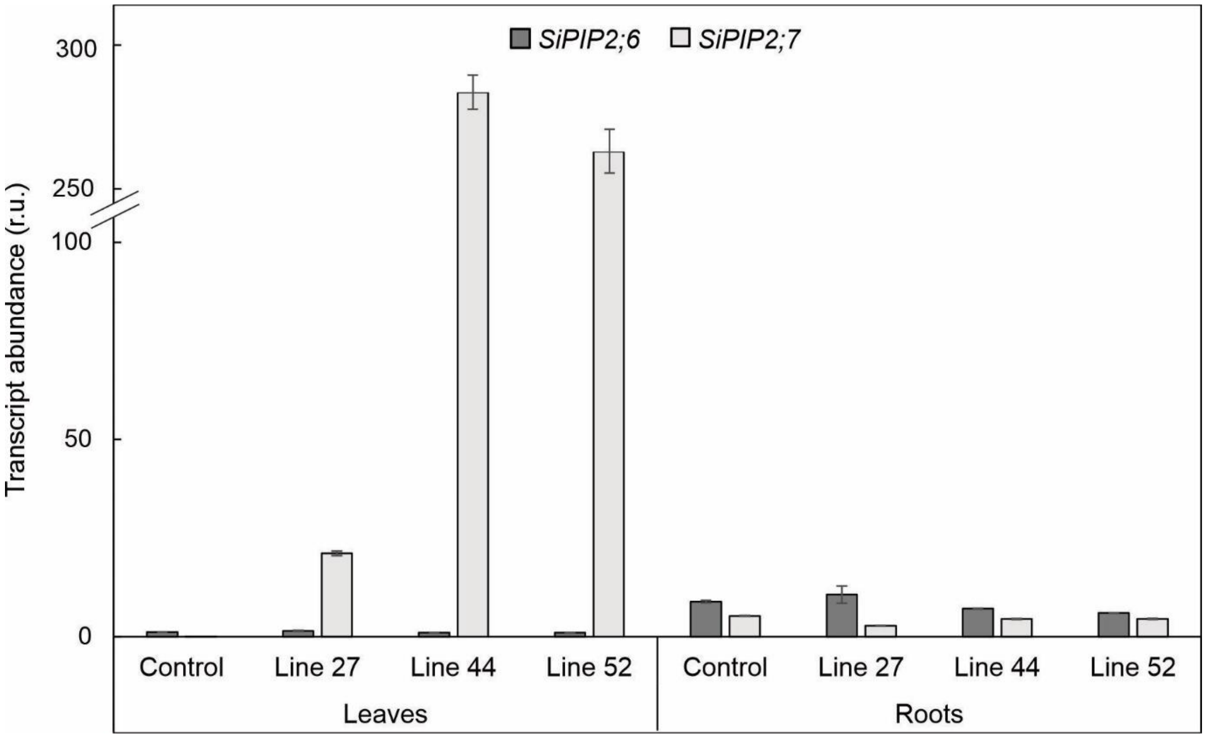


**Fig. S6**. qPCR analysis of *SiPIP2;6* and *SiPIP2;7* expression in leaves and roots of control and transgenic *S. viridis* plants. Azygous plants of line 44 were used as control.

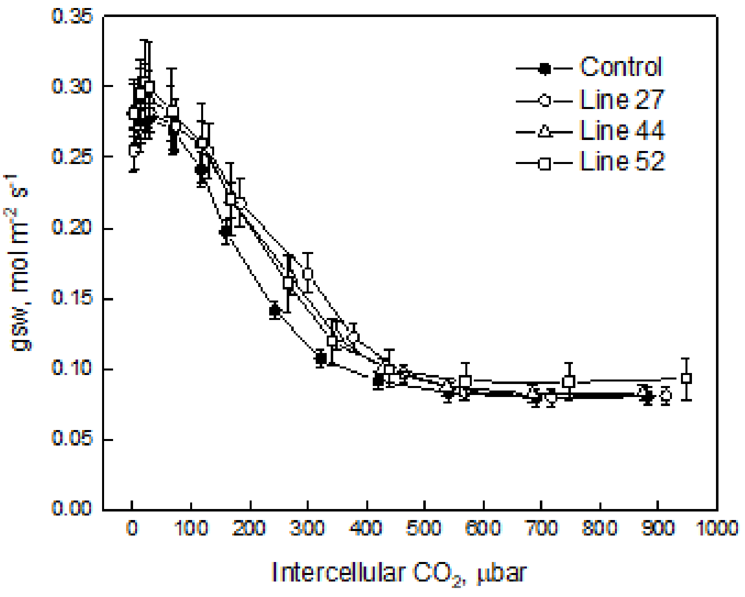


**Fig. S7**. Stomatal conductance to water vapor (gsw) measured at different intercellular $CO_2$ partial pressure. No significant differences were detected between the transgenic and control plants (One-way ANOVA, $\alpha = 0.05$).

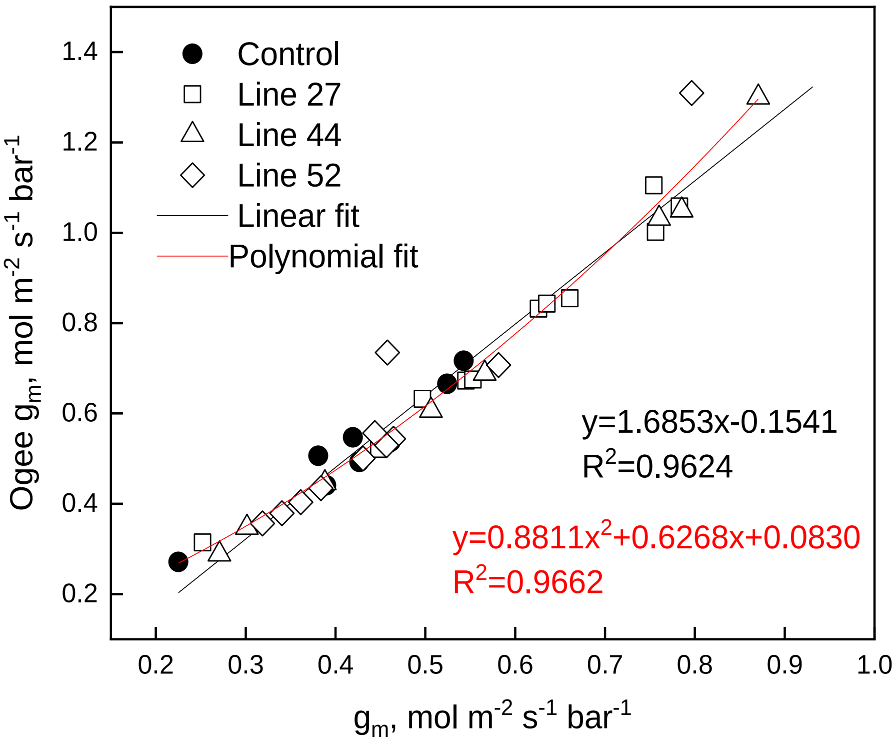


**Fig. S8**. Comparison of the $g_m$ estimated by oxygen isotope discrimination assuming full isotopic equilibrium [23] and calculations suggested by Ogée, et al. [34].

**Table S1**. Amino acid similarity of *S. italica* and *S. viridis* PIPs.

| *Setaria italica* | | | *Setaria viridis* | | |
|---|---|---|---|---|---|
| **Name** | **NCBI ID** | **AA length** | **Name** | **AA length** | **% identity** |
| SiPIP1;1 | Si010758m | 288 | SvPIP1;1 | 288 | 100 |
| SiPIP1;2 | Si017731m | 337 | SvPIP1;2 | 289 | 100 |
| SiPIP1;5 | Si017991m | 288 | SvPIP1;5 | 288 | 100 |
| SiPIP1;6 | Si007002m | 299 | SvPIP1;6 | 299 | 100 |
| SiPIP2;1 | Si030703m | 289 | SvPIP2;1 | 289 | 100 |
| SiPIP2;2 | Si039883m | 294 | SvPIP2;2 | 294 | 100 |
| SiPIP2;3 | Si036923m | 284 | SvPIP2;3 | 284 | 100 |
| SiPIP2;4 | Si017990m | 288 | SvPIP2;4 | 290 | 99.3 |
| SiPIP2;5 | Si010750m | 290 | SvPIP2;5 | 290 | 100 |
| SiPIP2;6 | Si030713m | 286 | SvPIP2;6 | 286 | 100 |
| SiPIP2;7 | Si030712m | 286 | SvPIP2;7 | 286 | 100 |
| SiPIP2;8 | Si030718m | 285 | SvPIP2;8 | 285 | 100 |

**Table S2**. Amino acid composition of *S. italica* PIPs at known substrate selectivity positions. Aquaporins have six transmembrane α-helices (H1-H6) joined by five loops (LA-LE). The monomeric channel is characterized by two NPA (Asn, Pro, Ala) motifs on loops A and E and along with the aromatic/arginine selectivity filter, these largely dictate the transport selectivity of substrates through the monomeric channels [6]. Froger's positions indicate additional residues that are predicted for substrate specificity [69].

| | | Gene Name | Gene Alias | Accession number | | NPA I motif | NPA II motif | ar/R | | | | | Froger's positions | | | | |
|---|---|---|---|---|---|---|---|---|---|---|---|---|---|---|---|---|---|
| | | *McGaughey et al, 2016* | *Azad et al, 2016* | Phytozome | NCBI | LB | LE | H2 | LC | H5 | LE1 | LE2 | P1 | P2 | P3 | P4 | P5 |
| PIP1 | Clade I | SiPIP1;1 | SiPIP1;2 | Seita.7G196700.1 | XP_004976483 | SGGHI**NPA**VT | GTGI**NPA**RSLG | F | G | H | T | R | Q | S | A | F | W |
| | | SiPIP1;2 | SiPIP1;1 | Seita.1G264900.1 | XP_004953388 | SGGHI**NPA**VT | GTGI**NPA**RSLG | F | G | H | T | R | Q | S | A | F | W |
| | | SiPIP1;5 | SiPIP1;3 | Seita.1G372300.1 | XP_004954451 | SGGHI**NPA**VT | GTGI**NPA**RSLG | F | G | H | T | R | Q | S | A | F | W |
| | | SiPIP1;6 | SiPIP1;4 | Seita.4G089800.1 | XP_004964964 | SGGHI**NPA**VT | GTGI**NPA**RSLG | F | G | H | T | R | V | S | A | F | W |
| PIP2 | Clade I | SiPIP2;1 | SiPIP2;5 | Seita.2G123000.1 | XP_004956113 | SGGHI**NPA**VT | GTGI**NPA**RSLG | F | G | H | T | R | Q | S | A | F | W |
| | | SiPIP2;4 | SiPIP2;4 | Seita.1G241900.1 | XP_004953172 | SGGHI**NPA**VT | GTGI**NPA**RSLG | F | G | H | T | R | Q | S | A | F | W |
| | | SiPIP2;5 | SiPIP2;3 | Seita.7G170200.1 | XP_004976254 | SGGHI**NPA**VT | GTGI**NPA**RSLG | F | G | H | T | R | Q | S | A | F | W |
| | | SiPIP2;6 | SiPIP2;2 | Seita.2G123200.1 | XP_004956115 | SGGHI**NPA**VT | GTGI**NPA**RSLG | F | G | H | T | R | Q | S | A | F | W |
| | | SiPIP2;7 | SiPIP2;1 | Seita.2G123300.1 | XP_004956116 | SGGHI**NPA**VT | GTGI**NPA**RSLG | F | G | H | T | R | Q | S | A | F | W |
| PIP2 | Clade II | SiPIP2;2 | SiPIP2;6 | Seita.9G219400.1 | XP_004986496 | SGGHI**NPA**VT | GTGI**NPA**RSFG | F | G | H | T | R | H | S | A | F | W |
| | | SiPIP2;3 | SiPIP2;7 | Seita.9G268100.1 | XP_012698174 | SGGHV**NPA**VT | GTGI**NPA**RSLG | F | G | H | T | R | H | S | A | F | W |
| | | SiPIP2;8 | SiPIP2;8 | Seita.2G291500.1 | XP_004957505 | SGGHI**NPA**VT | GTGI**NPA**RSLG | F | G | H | T | R | T | S | A | F | W |
| | | Substrate-specific signature sequences for $CO_2$ | | | | SGGHI**NPA**VT | GTGI**NPA**RSLG | F | - | H | T | R | Q/M | S | A | F | W |

**Table S3**. Primers used for qPCR

| Gene target | Forward | Reverse |
|---|---|---|
| *SvPIP2;6* | CAGAGCGGCTTCTACGCT | GCGTTGCGCTTGGGATCC |
| *SvPIP2;7* | TGTCCCTGGTGCGCGCAA | GAGAACGAAGGTGCCGACA |
| *Ubiquitin* | GATCTCCGCCCCAGCAAGAT | ATGCCCTCCTTGTCCTGGAT |
| *Elongation factor 1* | GCTGCAACAAGATGGATGCC | CCAGAGATTGGGACGAAGGC |
| *Tubulin* | CTAAAGCTCGCCACCCCTAC | GTCGGAGTTGAGCTGACCAG |
